## Supplementary Material for "Ubiquitination is a novel post-translational modification of VMP1 in autophagy of human tumor cells"

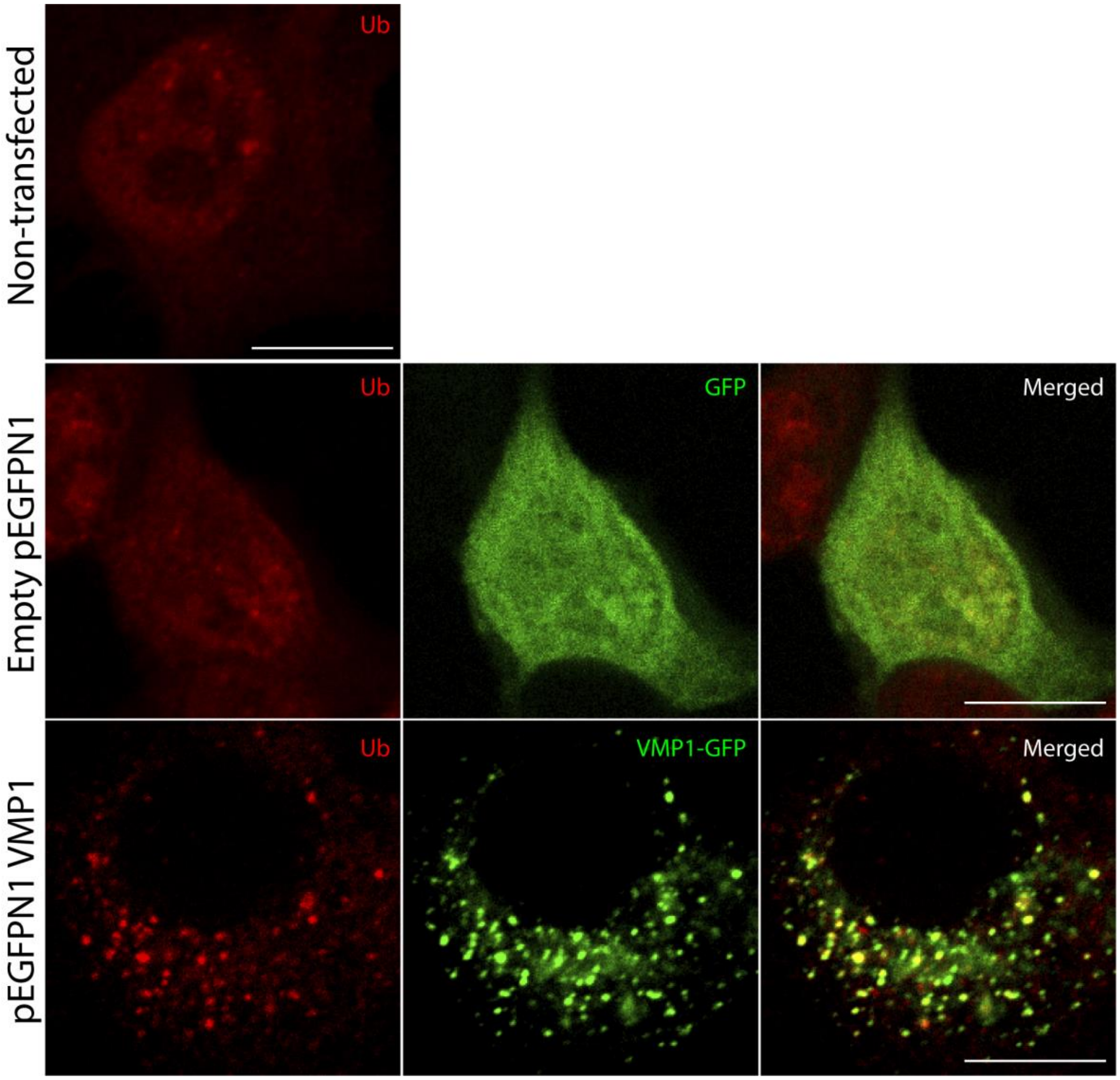


**Figure S1. Ubiquitin recruitment in PANC-1 cells.** PANC-1 cells expressing Empty pEGFPN1 or pEGFPN1 VMP1, or non-transfected PANC-1 cells, were immunolabeled with anti-ubiquitin. Scale bars: 10 μm.

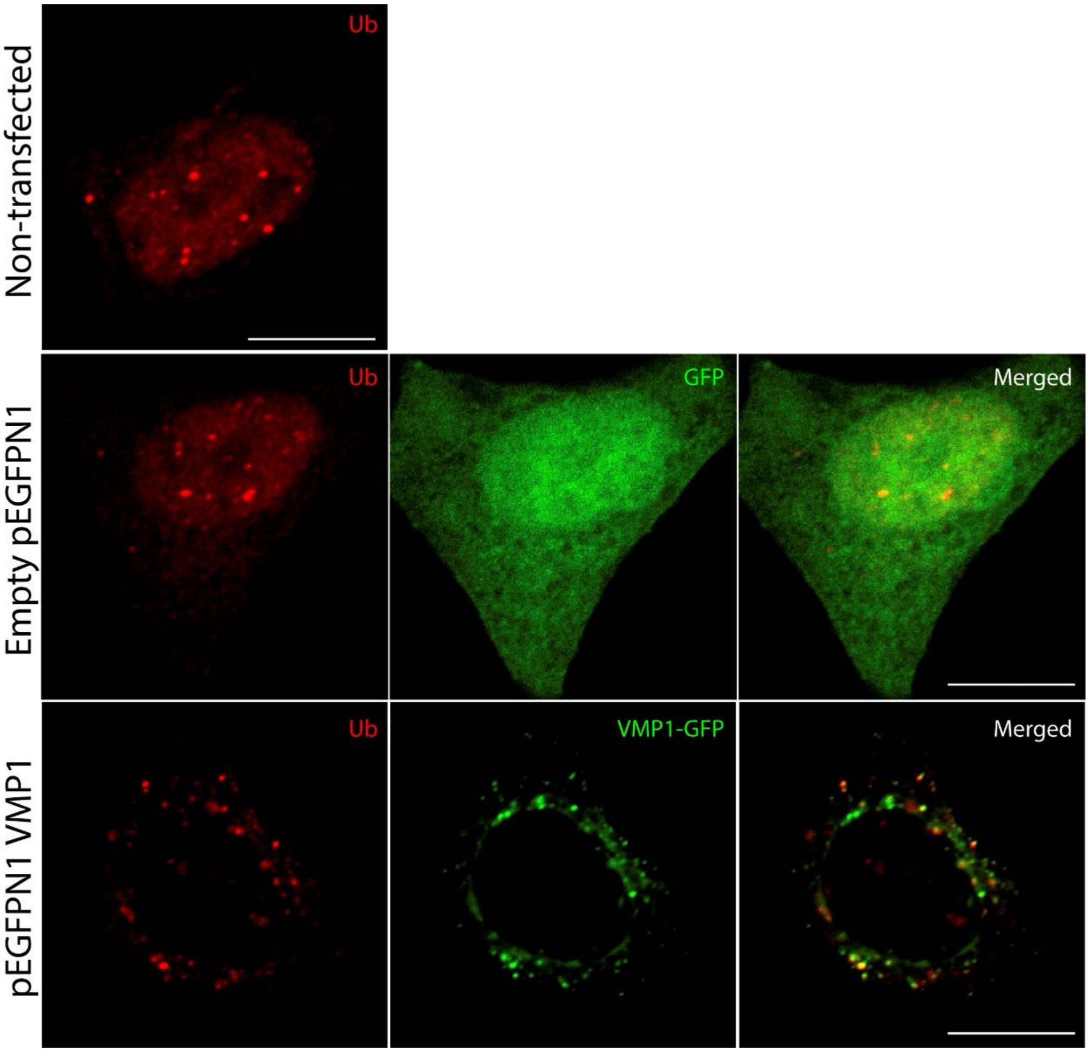


**Figure S2. Ubiquitin recruitment in MCF-7 cells.** MCF-7 cells expressing Empty pEGFPN1 or pEGFPN1 VMP1, or non-transfected MCF-7 cells, were immunolabeled with anti-ubiquitin. Scale bars: 10 μm.

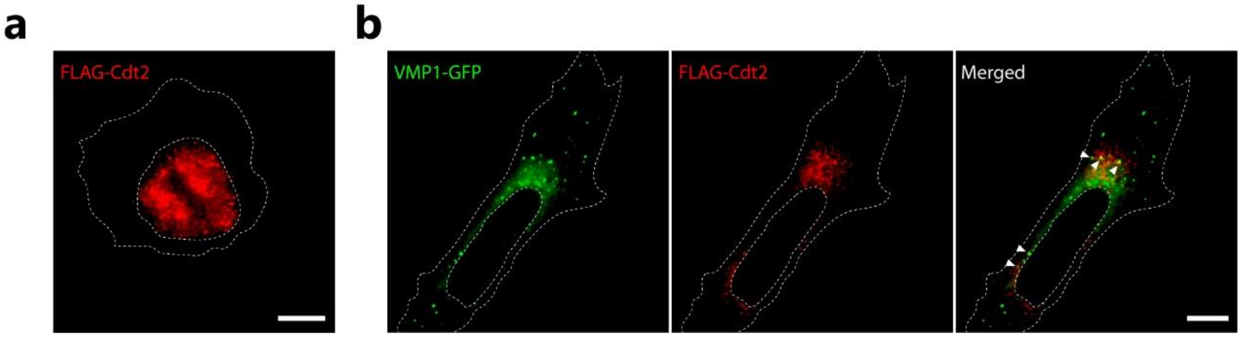

**Figure S3. FLAG-Cdt2 translocation.** (**a**) HeLa cells were transfected with FLAG-Cdt2 and immunolabeled with anti-FLAG. Scale Bar: 10 μm. (**b**) HeLa cells were transfected with VMP1-GFP and FLAG-Cdt2 and immunolabeled with anti-FLAG. Scale Bar: 10 μm. The images are representative of three independent experiments.
